# A Sparse Deep Learning Approach for Automatic Segmentation of Human Vasculature in Multispectral Optoacoustic Tomography

**DOI:** 10.1101/833251

**Authors:** Nikolaos-Kosmas Chlis, Angelos Karlas, Nikolina-Alexia Fasoula, Michael Kallmayer, Hans-Henning Eckstein, Fabian J Theis, Vasilis Ntziachristos, Carsten Marr

## Abstract

Multispectral Optoacoustic Tomography (MSOT) resolves oxy- (HbO_2_) and deoxy-hemoglobin (Hb) to perform vascular imaging. MSOT suffers from gradual signal attenuation with depth due to light-tissue interactions: an effect that hinders the precise manual segmentation of vessels. Furthermore, vascular assessment requires functional tests, which last several minutes and result in recording thousands of images. Here, we introduce a deep learning approach with a sparse UNET (S-UNET) for automatic vascular segmentation in MSOT images to avoid the rigorous and time-consuming manual segmentation. We evaluated the S-UNET on a test-set of 33 images, achieving a median DICE score of 0.88. Apart from high segmentation performance, our method based its decision on two wavelengths with physical meaning for the task-at-hand: 850 nm (peak absorption of oxy-hemoglobin) and 810 nm (isosbestic point of oxy-and deoxy-hemoglobin). Thus, our approach achieves precise data-driven vascular segmentation for automated vascular assessment and may boost MSOT further towards its clinical translation.

## 1 Introduction

The abundant presence of hemoglobin in the blood renders multispectral optoacoustic tomography (MSOT) an ideal technique for imaging vasculature [1–3]. By illuminating tissue at multiple different light wavelengths at the near infrared range (~680-980 nm), MSOT is capable of resolving several tissue chromophores, in particular oxy- (HbO_2_) and deoxy-hemoglobin (Hb), with a wide range of clinical applications, such as Crohn’s disease, systemic sclerosis, breast cancer, brown adipose tissue imaging and thyroid disease [4–8]. MSOT can provide precise structural visualizations of arteries and veins by recording multispectral data and resolving the different oxygenation states of human hemoglobin molecule. Moreover, the dynamic nature of the vascular system requires the acquisition not only of structural but also of functional data over multiple seconds or minutes to observe, for example, the vascular wall kinetics during the cardiac cycle or the arterial responses to stimuli such as the transient arterial occlusion or hyperthermia, which are valid descriptors of cardiovascular risk [9, 10]. The need to record multispectral data in order to extract molecular information and to perform longitudinal measurements over several minutes radically increases the number of recorded images and the data volume.

Both structural and functional vascular imaging requires the precise segmentation of the vascular lumen in several applications, such as the quantification of an atheromatous arterial stenosis, the detection of a venous thrombosis or the tracking of the arterial diameter over a 5-minute arterial occlusion challenge to quantify the degree of endothelial dysfunction. The segmentation of the vascular lumen is usually performed by expert physicians who manually draw the regions of interest (ROIs) on the recorded MSOT images. However, manual segmentation is a time-consuming process, in particular in the case of longitudinal recordings of several minutes and thus of hundreds or thousands of frames. Furthermore, because of the gradual light attenuation due to scattering and absorption when propagating in living tissue, the vascular lumen shows an inhomogeneous and fainting intensity profile with increasing depth, making its manual delineation a challenging process. But even in routine imaging diagnostics, a reliable automated segmentation method can be beneficial by aiding the clinician in performing the same task much faster. Deep learning has been recently shown to be very effective in computer vision tasks [11, 12] and segmentation in particular [13—15]. As such, deep learning has been successfully applied to clinical diagnostics [16–18], with medical image segmentation applications including prostate [19], retinal disease [20], brain [21, 22] and cervical cell segmentation [23]. Surveys of deep learning applications for medical imaging can be found in [24, 25].

We present herein a pilot study to achieve automated vascular segmentation in clinical raw MSOT images via a deep learning approach, based on an extension of the UNET architecture originally introduced in [15] that is specifically tailored for multispectral optoacoustic data. The proposed Sparse UNET (S-UNET) allows for automated segmentation of vascular ROIs in clinical MSOT images, while simultaneously identifying which of the employed illumination wavelengths are relevant to the specific task. This way we aim at radically reducing the time needed for vascular segmentation in longitudinal scans as well as the number of illumination wavelengths for future task-specific scans, facilitating this way the data analysis, increasing the time resolution and reducing the data volume.

## 2 Methods

### 2.1 Network Architecture

The proposed Sparse-UNET (S-UNET) is based on the fully convolutional architecture of the original UNET [5], with the added capability of sparse wavelength selection. The goal of S-UNET is to transform each input image with dimensions 400×400×28 (Height × Width × Wavelengths) into a 400×400 probability map ***p*** that corresponds to a ground truth segmentation mask, while simultaneously assigning a weight *w*_*c*_ (= wavelength importance) to each of the 28 illumination wavelengths (from 700 to 970 nm at steps of 10 nm). The ground truth segmentation mask ***y*** is a binary image (each pixel is either 0 or 1), extracted from the recorded MSOT image in consensus between two clinical MSOT experts. To arrive at a predicted segmentation mask, the resulting S-UNET probability map ***p*** is discretized by thresholding at 0.5: pixels with probabilities less than 0.5 are set to 0, while the rest are set to 1.

In order to perform wavelength selection, the first layer of the S-UNET corresponds to a 1×1 2D convolution of a single filter and no bias. Given each 400×400×28 input image stack, the first layer essentially performs a linear combination of the 28 wavelengths, resulting in a 400×400×1 image that is forward-propagated to the rest of the network. In this manner, each wavelength is assigned a unique scalar weight. Moreover, to ensure sparsity of wavelength selection we add L1 regularization [26] on the wavelength weights. Regularization does not necessarily result in an interpretable model. To ensure interpretability of wavelength selection we force the weights of the first layer to be non-negative. As such, there is no possibility to have irrelevant wavelengths of similar wavelengths cancelling each other out with weights of similar, potentially high, magnitude and opposite signs. Taken together, the two constraints of L1 regularization and non-negative weights ensure that only few relevant wavelengths will be assigned with positive weights, while all other non-relevant wavelengths will be set to zero and will effectively be excluded from the model. After wavelength selection, we add a batch normalization [27] layer between every convolution layer and its respective activation function. The S-UNET architecture employed is visualized in Fig. 1.

**Fig. 1.**
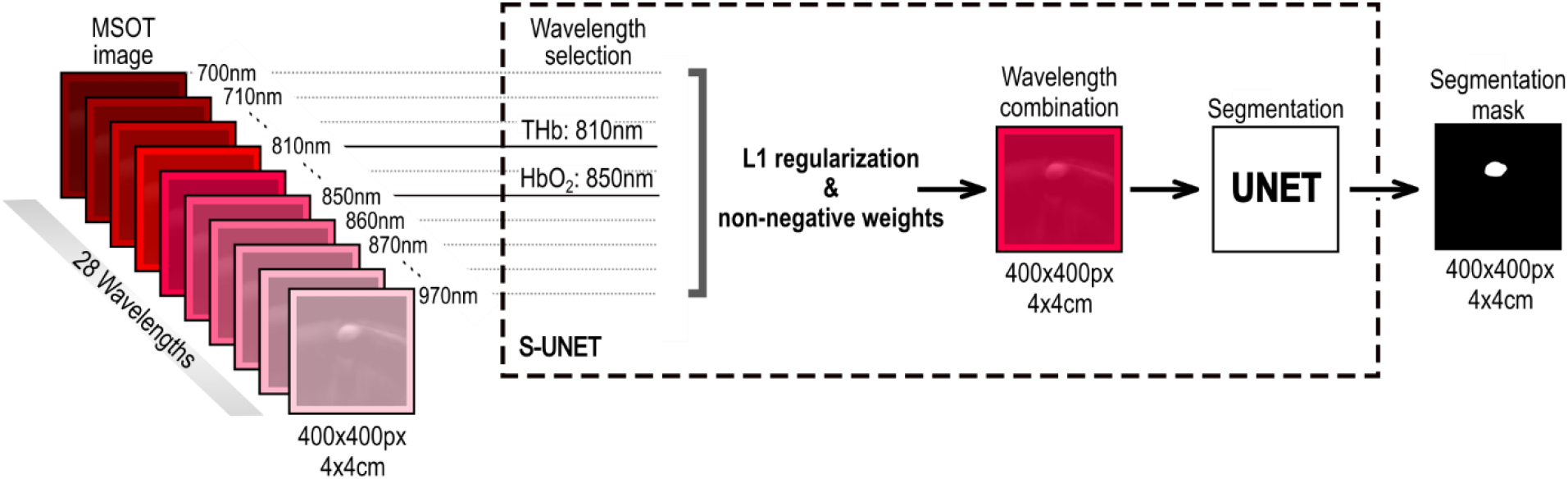
The S-UNET identifies important illumination wavelengths in MSOT images while learning to predict segmentation masks of human blood vessels. Each wavelength is weighted by a corresponding non-negative weight and all weighted wavelengths are combined before being inserted as input into a UNET architecture. Sparsity of wavelength selection is enforced by L1 regularization on the non-negative wavelength weights and the weights themselves are learned through standard back-propagation, along with the rest of the UNET parameters.

### 2.2 Training and Data Augmentation

The original dataset of 164 raw MSOT images is randomly split into training, validation and test sets of 98, 33 and 33 images, respectively.

Each raw MSOT image corresponds to spatial dimensions of 400×400 pixels (which corresponds to 4×4 cm) and 28 wavelengths. Each wavelength is normalized to values between 0 and 1 separately, as part of pre-processing. We train the model on a training subset of the data using Adam [28] while evaluating model performance on a validation set. The model is trained for a maximum number of 200 epochs, or until model performance on the validation set has not improved for 20 consecutive epochs (early stopping). The instance of the model that achieved the best performance on the validation set is saved as the final model. We keep a separate test set that is hidden from the model during training.

The model is trained using a batch size of 4 images and data augmentation is performed on-the-fly on each image in every batch to increase model performance. Data augmentation includes flipping the x axis and rotating the image in a random angle from 5 to 15 degrees. Each of the two augmentation schemes has a 50% probability of being performed on any given image. According to our experiments more aggressive augmentation hinders model performance on the given task. The last layer of the model corresponds to a pixel-wise binary classification problem of computing a probability map of the predicted segmentation mask. The model’s loss corresponds to the loss of the 400×400 binary classification tasks. Thus, the total binary cross entropy loss function *L*, is used to train the model:

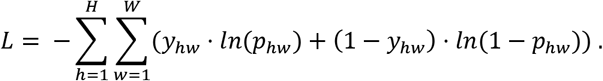

Here, *H* and *W*correspond to the image height and width in pixels (each being 400), *y*_*hw*_ ∊ {0,1} corresponds to the ground truth segmentation class, *p*_*hw*_ ∈ [0,1] corresponds to the predicted class probability for the corresponding pixel in position (*h*, *w)* and *ln* is the natural logarithm.

## 3 Experiments

### 3.1 Data Acquisition

In this pilot study we scanned six (n=6) healthy volunteers (3 men, 3 women, age 30 ± 5.44 years). All healthy volunteers consented to participate in this study in full accordance with the work safety regulations of the Helmholtz Center Munich (Neuherberg, Germany). The radial artery, the brachial artery, the dorsal artery of the foot, as well as the cephalic vein, the radial veins and the dorsal vein of the foot were scanned by means of a clinical hand-held MSOT/Ultrasound system (iThera Medical GmbH, Munich, Germany). All subjects were asked to consume no food or caffeine for 8 hours before the examination, which was conducted in a quiet dark room with normal temperature of 25°C. Each scan lasted for 5-10 seconds. The system used was equipped with a near-infrared laser for achieving optimal penetration depth in tissue (3-4 cm) even with low illumination energy (~15 mJ per pulse). For multispectral data recording we used 28 wavelengths (700:10:980 nm). Tissue was illuminated by short light pulses (~10 ns) at a frame rate of 25fps. The ultrasound detection was performed by 256 ultrasound sensors with a central frequency of 4 MHz which covered an angle of 145° and was mounted on the hand-held scanning probe. Acquired ultrasound signals for each illumination pulse were reconstructed into a tomographic image using a model-based reconstruction algorithm [29]. For each MSOT image a co-registered ultrasound image was recorded. The segmentation of the scanned arteries and veins was manually performed on the appropriate MSOT frame by simultaneous view of the co-registered ultrasound image. We decided to segment the blood vessels directly on the MSOT frames because of better contrast, compared to ultrasound, provided by the high light absorption of hemoglobin at the near-infrared illumination range. The appropriate frame for vein segmentation was the frame corresponding to the 750 nm illumination wavelength were the absorption of Hb is clearly higher than that of HbO_2_. The appropriate frame for artery segmentation was the frame corresponding to the 850 nm illumination wavelength were the absorption of HbO_2_ is clearly higher than that of Hb. Manual segmentation was conducted in consensus of two clinicians with experience in MSOT and clinical ultrasound imaging.

### 3.2 Model Comparison and Wavelength Selection

We compared three segmentation methods on the MSOT images described above: the proposed S-UNET as well as two differently sized variants of a standard UNET to the segmentation task. The S-UNET corresponds to the architecture described in the previous section. The wavelength selection layer is followed by a downsized UNET where every convolutional layer corresponds to 1/8 of filters compared to the architecture in [15]. Two variants of the UNET were applied: one with the same number of filters as in [15] (‘original’) and a variant with 1/8 of filters (‘downsized’). Additionally, a batch normalization layer was inserted between every convolutional layer and its corresponding activation function in both UNET variants. Training was performed as described in the previous section. The results of all segmentation methods were compared to the binary ground truth segmentation mask, which was manually generated from expert clinicians on the recorded MSOT images under co-registered ultrasound guidance (see Methods). Model comparison is based on the Dice coefficient [22] defined as:

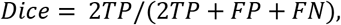

where TP, FP and FN correspond to true positive, false positive and false negative classified pixels: A TP pixel is a correctly classified foreground pixel, a FP pixel is a background pixel falsely classified as foreground, and a FN pixel corresponds to a foreground pixel that was incorrectly classified as background by the model. The Dice coefficient is well-suited to tackle the class imbalance inherent to the segmentation task [30], where more than 99% of the pixels in our dataset are background pixels. As such, it is preferred for model assessment compared to the standard cross entropy used to train the model. The Dice coefficient lies between 0 and 1, with higher values being better since they correspond to larger overlap between the ground truth and predicted segmentation masks.

### 3.3 Wavelength Selection

Wavelength selection was performed by the first layer of the S-UNET (see Methods). However, since feature selection is an inherently noisy process [31, 32] it is good practice to average a number of models [33] in order to obtain a smoothed version of wavelength importance. We thus train 100 different instances of the S-UNET and aggregate their results for the tasks of segmentation, as well as for wavelength selection. In the case of segmentation, we average the probability maps of all models before discretizing in order to obtain the binary segmentation mask.

### 3.4 Results

The performance of all three segmentation models is comparable as reported by the Dice coefficient (see Table 1). The original UNET with over 30 million parameters is potentially slightly overfitting the training dataset while the downsized UNET, as well as the S-UNET achieve very similar segmentation results with roughly half a million parameters. The downsized UNET achieves slightly higher Dice scores on average (0.90±0.08) than the S-UNET ensemble (0.86±0.11), but the difference is not statistically significant given the test set size of 33 images (p-value = 0.37, two-sample Wilcoxon rank-sum test). This similarity in performance is to be expected since both methods correspond to a similar number of parameters. However, this also suggests that the added sparsity of wavelength selection does not affect, at least not significantly, the quality of the generated segmentation masks in the case of S-UNET.

**Table 1.** Model Performance. Dice results correspond to mean±std.

| Model | Test Set Dice | Parameters | Wavelength Selection |
| --- | --- | --- | --- |
| UNET (original) | 0.75±0.28 | 31,416,897 | No |
| UNET (downsized) | 0.90±0.08 | 495,881 | No |
| S-UNET | 0.86±0.11 | 493,965 | Yes |

The advantage of the S-UNET approach over the two UNET approaches is clearly its interpretability of results due to the embedded wavelength selection. As visualized in Fig. 2, out of the 28 input wavelengths the model has identified two as being the most important in a purely data-driven manner. These wavelengths correspond to the maximum of the absorption spectra of total blood volume (HbO_2_ and Hb) at 810nm and HbO_2_ at 850nm. Both of these identified wavelengths are thus meaningful since they mark the presence of blood in the detected image regions. The segmentation results of S-UNET on an exemplary set of images are visualized in Fig. 3. Interestingly, our approach is able to discriminate blood vessels from similar objects probably by exploiting the wavelength information.

**Fig. 2.**
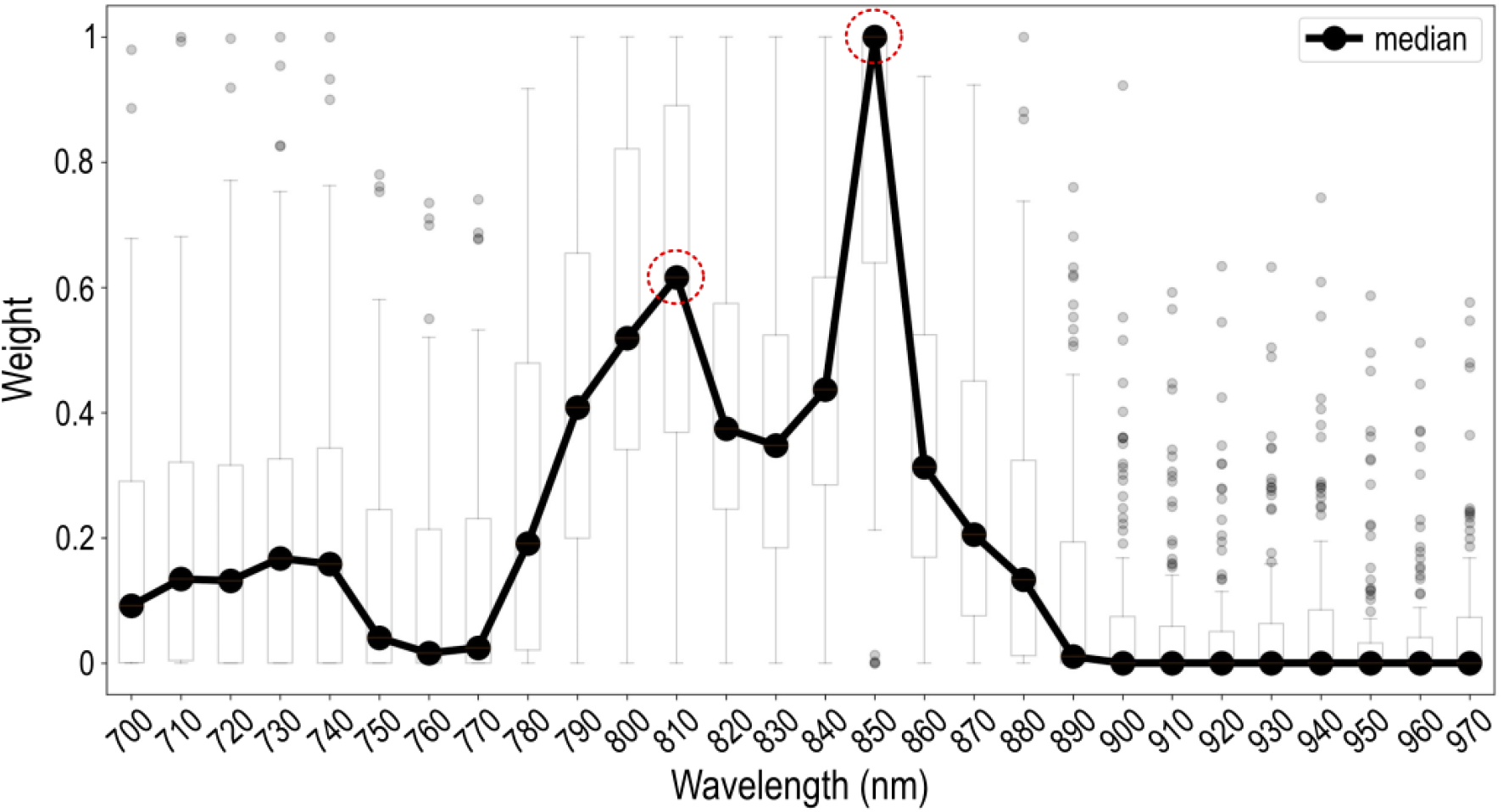
The S-UNET identifies wavelengths relevant to the segmentation task. Each boxplot (the box’s edges correspond to quartiles 1 and 3 while whiskers extend to ±1.5 times the interquartile range) corresponds to the weights assigned by the ensemble of 100 S-UNET instances to each wavelength. Averaging results is necessary since feature selection is an inherently noisy process. According to the median weight of each wavelength, the two most important wavelengths are 850 nm and 810 nm, corresponding to the maximum absorption of HbO_2_ and total hemoglobin (THb), respectively.

**Fig. 3.**
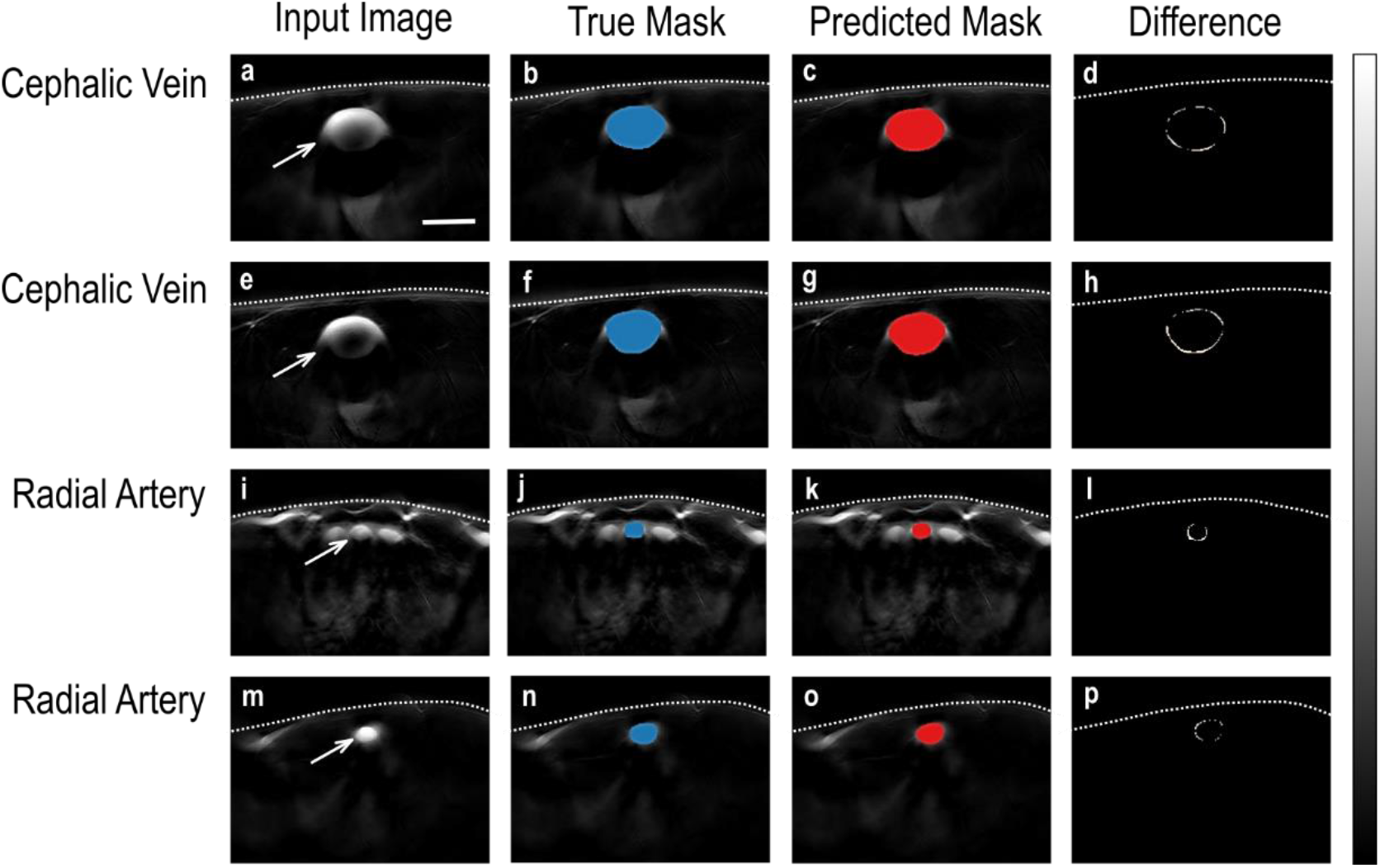
The S-UNET successfully segments human vasculature from MSOT images. Each row corresponds to a different image of the test set. The first column (images a, e, i, m) shows the 850nm channel of the MSOT image. The second (images b, f, j, n) and third columns (c, g, k, o) show the ground truth (true mask, blue) and predicted segmentation masks (red), respectively, visualized on top of the input image. The true segmentation mask is identified by expert physicians, while the S-UNET predicted segmentation mask corresponds to the output of the S-UNET ensemble. The fourth column (images d, h, l, p) corresponds to the absolute difference between the true and predicted binary segmentation masks and is equivalent to the logical operation of XOR (exclusive or). The predicted masks almost completely overlap with the ground truth segmentation. The S-UNET is successful even in the last two cases (rows) where the mask is relatively small and located in an area where similar bright spots are present. The white dashed line represents the skin surface. The white arrows point to the blood vessel of interest. The scale bar is 5 mm. The gray color bar ranges from 0 to 1 and corresponds to the normalized intensity of each image (columns 1-3) or the difference of the true and predicted segmentation masks (column 4).

## 4 Discussion

In this work we applied a deep learning approach based on an adapted S-UNET to perform automated vascular segmentation in clinical MSOT images. Our model successfully segments blood vessels (arteries and veins) and its performance is comparable to a standard UNET of similar model size. Furthermore, our model is capable of selecting the illumination wavelengths that are most important for the segmentation task at hand in a purely data driven manner. Our results show that among the 28 illumination wavelengths used for data acquisition, two wavelengths are associated with the light absorption of hemoglobin at the near-infrared range of illumination (700 to 970 nm). These correspond to 810 nm, which is the isosbestic point of HbO_2_ and Hb and reflects the absorption of total hemoglobin or else the total blood within the vasculature and 850nm, which is the point where HbO_2_ absorbs significantly more than Hb and reflects the arterial blood.

Our approach achieves accurate automated segmentation of both arteries and veins on raw clinical MSOT data. Apart from facilitating the segmentation process, which is time-costly for longitudinal scans of several minutes during functional vascular testing, it may help tackling a significant limitation of optical and optoacoustic imaging: the attenuation of light due to scattering and absorption when propagating in living tissue. This effect causes a gradual attenuation of measured intensity in the vascular lumen with increasing depth rendering the partial or even the total visualization and thus the accurate segmentation of it a real challenge even for clinicians with extensive MSOT experience. In the current study, we scanned blood vessels where this effect was apparent (e.g. Fig. 3a) but not to an extent that would jeopardize the accurate manual segmentation of the vascular lumen directly on the MSOT images under ultrasound guidance. Thus, future studies are required to further investigate the efficacy of deep learning approaches in automatically detecting and segmenting vessels with clinical interest (e.g. the carotid artery) deep in tissue in clinical MSOT data.

In this study, we preferred to work on raw MSOT data. However, the discrete spectral difference of HbO2 and Hb at the near-infrared range as well as the strong presence of HbO2 in arteries and Hb in veins would allow for the direct spectral unmixing of HbO2 and Hb in the MSOT data and thus for direct vascular segmentation. Nevertheless, spectrally unmixed data suffer from errors related to imaging depth and motion, either exogenous (e.g. operator’s hand and random patient movement) or endogenous (e.g. arterial pulsation or breathing).

Regarding motion-related errors, the dynamic character of the vascular system introduces significant inaccuracy when it comes to spectral unmixing results, especially when illuminating at multiple different wavelengths (e.g. 28) to achieve high spectral quality. For example, the recording of a multispectral stack of 28 wavelengths at a frame rate of 25 Hz takes more than one second. Considering that the cardiac cycle of a normal individual with a heart rate of 70-75 Hz is approximately 0.8 sec, the use of 28 wavelengths renders the spectrally unmixed data vulnerable to errors due to arterial wall motion, especially in the periphery of the vascular lumen, potentially degrading the precision of vascular segmentation when performed by means of direct spectral unmixing.

Moreover, multispectral optoacoustic imaging at increased tissue depths (> 1cm), where normally the blood vessels lie, renders the spectral unmixing output vulnerable to the spectral coloring effect: the random absorption of each illumination wavelength before reaching the HbO2 or the Hb of the vascular lumen according to the light absorption properties of the set of tissues covering them (e.g. skin, subcutaneous fat, muscle). Thus, usual linear spectral unmixing methods fail to unmix the absorbers of interest (e.g. HbO2 and Hb) at increasing depths since the measured spectra have been colored and thus deflected from the known absorption spectra, as measured in the lab. For the abovementioned reasons, we decided to work on the recorded raw MSOT data.

Our model showed that the decision for segmenting the vasculature was mainly based on two near-infrared wavelengths: the 810 nm where HbO_2_ and Hb absorb light to the same extent and the 850 nm where the light absorption of HbO_2_ is significantly higher than that of Hb. Our results provide evidence for effective and task-specific wavelength selection via the suggested deep learning model for accurate segmentation of blood vessels in clinical MSOT data. Apart from increasing the time resolution by skipping a number of unnecessary illumination wavelengths and decreasing the data volume, the effective wavelength selection may be used for indirect spectral characterization of more complex tissues or even homogeneous tissues at high depths by identifying the wavelengths critical for achieving their segmentation. This approach may help overcoming the limitations introduced by the spectral coloring effect and thus providing a blind or data-driven spectral unmixing with great implications for clinical MSOT imaging. Our method may be used for segmenting and characterizing tissues with clinical relevance (e.g. the subcutaneous fat or the atherosclerotic plaques which contain lipids, the skeletal muscle which contains water) or even the detection and distribution mapping of injected contrast agents targeting specific molecules involved in the pathophysiology of a disease.

To the best of our knowledge, while deep learning has been used before in the context of optoacoustic imaging data [34, 35], this is the first time where a deep learning method is applied to clinical MSOT data. Our approach has significant implications for future MSOT applications with clinical relevance, such as the automated segmentation of more complex soft tissues (e.g. muscle, fat, atherosclerotic plaques) and foreseeable for more accurate diagnosis of vascular disease.

## Acknowledgements

N.K.C. acknowledges support from the Graduate School of Quantitative Biosciences Munich (QBM). F.J.T. acknowledges financial support by the Graduate School QBM, by the Helmholtz Association (Incubator grant sparse2big, grant # ZT-I-0007), by the BMBF (grant# 01IS18036A and grant# 01IS18053A) and by the Chan Zuckerberg Initiative DAF (advised fund of Silicon Valley Community Foundation, 182835). C.M acknowledges support from the BMBF (e:Med grant MicMode-I2T). This project has received funding from the European Research Council (ERC) under the European Union’s Horizon 2020 research and innovation program under grant agreement No 694968 (PREMSOT) and was supported by the DZHK (German Centre for Cardiovascular Research) and by the Helmholtz Zentrum München, funding program “Physician Scientists for Groundbreaking Projects”.

## References

1. A. Karlas, N. A. Fasoula, K. Paul-Yuan, J. Reber, M. Kallmayer, D. Bozhko, M. Seeger, H. H. Eckstein, M. Wildgruber, and V. Ntziachristos, “Cardiovascular optoacoustics: From mice to men - A review,” Photoacoustics 14, 19–30 (2019).

2. A. Karlas, J. Reber, G. Diot, D. Bozhko, M. Anastasopoulou, T. Ibrahim, M. Schwaiger, F. Hyafil, and V. Ntziachristos, “Flow-mediated dilatation test using optoacoustic imaging: a proof-of-concept,” Biomed Optics Express 8, 3395–3403 (2017).

3. M. Masthoff, A. Helfen, J. Claussen, A. Karlas, N. A. Markwardt, V. Ntziachristos, M. Eisenblatter, and M. Wildgruber, “Use of Multispectral Optoacoustic Tomography to Diagnose Vascular Malformations,“ JAMA dermatology (2018).

4. W. Roll, N. A. Markwardt, M. Masthoff, A. Helfen, J. Claussen, M. Eisenblatter, A. Hasenbach, S. Hermann, A. Karlas, M. Wildgruber, V. Ntziachristos, and M. Schafers, “Multispectral optoacoustic tomography of benign and malignant thyroid disorders - a pilot study,“ Journal of nuclear medicine : official publication, Society of Nuclear Medicines (2019).

5. J. Reber, M. Willershäuser, A. Karlas, K. Paul-Yuan, G. Diot, D. Franz, T. Fromme, S. V. Ovsepian, N. Bézière, E. Dubikovskaya, D. C. Karampinos, C. Holzapfel, H. Hauner, M. Klingenspor, and V. Ntziachristos, “Non-invasive Measurement of Brown Fat Metabolism Based on Optoacoustic Imaging of Hemoglobin Gradients,” Cell Metabolism 27, 689–701.e684 (2018).

6. M. Masthoff, A. Helfen, J. Claussen, W. Roll, A. Karlas, H. Becker, G. Gabriels, J. Riess, W. Heindel, M. Schafers, V. Ntziachristos, M. Eisenblatter, U. Gerth, and M. Wildgruber, “Multispectral optoacoustic tomography of systemic sclerosis,” J Biophotonics 11, e201800155 (2018).

7. G. Diot, S. Metz, A. Noske, E. Liapis, B. Schroeder, S. V. Ovsepian, R. Meier, E. Rummeny, and V. Ntziachristos, “Multispectral Optoacoustic Tomography (MSOT) of Human Breast Cancer,” Clinical cancer research : an official journal of the American Association for Cancer Research 23, 6912–6922 (2017).

8. F. Knieling, C. Neufert, A. Hartmann, J. Claussen, A. Urich, C. Egger, M. Vetter, S. Fischer, L. Pfeifer, A. Hagel, C. Kielisch, R. S. Gortz, D. Wildner, M. Engel, J. Rother, W. Uter, J. Siebler, R. Atreya, W. Rascher, D. Strobel, M. F. Neurath, and M. J. Waldner, “Multispectral Optoacoustic Tomography for Assessment of Crohn’s Disease Activity,” N Engl J Med 376, 1292–1294 (2017).

9. D. J. Green, H. Jones, D. Thijssen, N. T. Cable, and G. Atkinson, “Flow-Mediated Dilation and Cardiovascular Event Prediction,” (2011).

10. S. Agarwal, J. Allen, A. Murray, and I. Purcell, “Comparative reproducibility of dermal microvascular blood flow changes in response to acetylcholine iontophoresis, hyperthermia and reactive hyperaemia,” Physiological measurement 31, 1–11 (2010).

11. A. Krizhevsky, I. Sutskever, and G. E. Hinton, “ImageNet Classification with Deep Convolutional Neural Networks,” in Advances in Neural Information Processing Systems 25, F. Pereira, C. J. C. Burges, L. Bottou, and K. Q. Weinberger, eds. (Curran Associates, Inc., 2012), pp. 1097–1105.

12. K. He, X. Zhang, S. Ren, and J. Sun, “Deep Residual Learning for Image Recognition,” (2015).

13. V. Badrinarayanan, A. Kendall, and R. Cipolla, “SegNet: A Deep Convolutional Encoder-Decoder Architecture for Image Segmentation,” IEEE Transactions on Pattern Analysis and Machine Intelligence 39, 2481–2495 (2017).

14. J. Long, E. Shelhamer, and T. Darrell, “Fully convolutional networks for semantic segmentation,” in 2015 IEEE Conference on Computer Vision and Pattern Recognition (CVPR), 2015), 3431–3440.

15. O. Ronneberger, P. Fischer, and T. Brox, “U-Net: Convolutional Networks for Biomedical Image Segmentation,” in Medical Image Computing and Computer-Assisted Intervention – MICCAI 2015, (Springer International Publishing, 2015), 234–241.

16. U. R. Acharya, S. L. Oh, Y. Hagiwara, J. H. Tan, M. Adam, A. Gertych, and R. S. Tan, “A deep convolutional neural network model to classify heartbeats,” Computers in Biology and Medicine 89, 389–396 (2017).

17. A. Esteva, B. Kuprel, R. A. Novoa, J. Ko, S. M. Swetter, H. M. Blau, and S. Thrun, “Dermatologist-level classification of skin cancer with deep neural networks,” Nature 542, 115–118 (2017).

18. P. Rajpurkar, J. Irvin, R. L. Ball, K. Zhu, B. Yang, H. Mehta, T. Duan, D. Ding, A. Bagul, C. P. Langlotz, B. N. Patel, K. W. Yeom, K. Shpanskaya, F. G. Blankenberg, J. Seekins, T. J. Amrhein, D. A. Mong, S. S. Halabi, E. J. Zucker, A. Y. Ng, and M. P. Lungren, “Deep learning for chest radiograph diagnosis: A retrospective comparison of the CheXNeXt algorithm to practicing radiologists,” PLoS medicine 15, e1002686 (2018).

19. T. Brosch, J. Peters, A. Groth, T. Stehle, and J. Weese, “Deep Learning-Based Boundary Detection for Model-Based Segmentation with Application to MR Prostate Segmentation,” in Lecture Notes in Computer Science (Springer International Publishing, 2018), 515–522.

20. J. D. Fauw, J. R. Ledsam, B. Romera-Paredes, S. Nikolov, N. Tomasev, S. Blackwell, H. Askham, X. Glorot, B. O’Donoghue, D. Visentin, G. v. d. Driessche, B. Lakshminarayanan, C. Meyer, F. Mackinder, S. Bouton, K. Ayoub, R. Chopra, D. King, A. Karthikesalingam, C. O. Hughes, R. Raine, J. Hughes, D. A. Sim, C. Egan, A. Tufail, H. Montgomery, D. Hassabis, G. Rees, T. Back, P. T. Khaw, M. Suleyman, J. Cornebise, P. A. Keane, and O. Ronneberger, “Clinically applicable deep learning for diagnosis and referral in retinal disease,” Nature Medicine 24, 1342 (2018).

21. K. Kamnitsas, C. Ledig, V. F. J. Newcombe, J. P. Simpson, A. D. Kane, D. K. Menon, D. Rueckert, and B. Glocker, “Efficient multi-scale 3D CNN with fully connected CRF for accurate brain lesion segmentation,” Medical Image Analysis 36, 61–78 (2017).

22. S. Pereira, A. Pinto, V. Alves, and C. A. Silva, “Brain Tumor Segmentation Using Convolutional Neural Networks in MRI Images,” IEEE transactions on medical imaging 35, 1240–1251 (2016).

23. Y. Song, E. Tan, X. Jiang, J. Cheng, D. Ni, S. Chen, B. Lei, and T. Wang, “Accurate Cervical Cell Segmentation from Overlapping Clumps in Pap Smear Images,” IEEE transactions on medical imaging 36, 288–300 (2017).

24. J. Ker, L. Wang, J. Rao, and T. Lim, “Deep Learning Applications in Medical Image Analysis,” IEEE Access 6, 9375–9389 (2018).

25. G. Litjens, T. Kooi, B. E. Bejnordi, A. A. A. Setio, F. Ciompi, M. Ghafoorian, J. A. W. M. van der Laak, B. van Ginneken, and C. I. Sánchez, “A survey on deep learning in medical image analysis,” Medical Image Analysis 42, 60–88 (2017).

26. R. Tibshirani, “Regression Shrinkage and Selection via the Lasso,” Journal of the Royal Statistical Society. Series B (Methodological) 58, 267–288 (1996).

27. S. Ioffe and C. Szegedy, “Batch Normalization: Accelerating Deep Network Training by Reducing Internal Covariate Shift,” in International Conference on Machine Learning, 2015), 448–456.

28. D. P. Kingma and J. Ba, “Adam: A Method for Stochastic Optimization,” arXiv:1412.6980 [cs] (2014).

29. A. Rosenthal, V. Ntziachristos, and D. Razansky, “Model-based optoacoustic inversion with arbitrary-shape detectors,” Medical physics 38, 4285–4295 (2011).

30. H. Kervadec, J. Bouchtiba, C. Desrosiers, E. Granger, J. Dolz, and I. B. Ayed, “Boundary loss for highly unbalanced segmentation,” in International Conference on Medical Imaging with Deep Learning, 2019), 285–296.

31. A. Kalousis, J. Prados, and M. Hilario, “Stability of feature selection algorithms,” in Fifth IEEE International Conference on Data Mining (ICDM’05), 2005), 8-pp.-.

32. S. Nogueira, K. Sechidis, and G. Brown, “On the Stability of Feature Selection Algorithms,” Journal of Machine Learning Research 18, 1–54 (2018).

33. N. Chlis, E. S. Bei, and M. Zervakis, “Introducing a Stable Bootstrap Validation Framework for Reliable Genomic Signature Extraction,” IEEE/ACM Transactions on Computational Biology and Bioinformatics 15, 181–190 (2018).

34. D. Allman, A. Reiter, and M. A. L. Bell, “Photoacoustic Source Detection and Reflection Artifact Removal Enabled by Deep Learning,” IEEE transactions on medical imaging 37, 1464–1477 (2018).

35. S. Antholzer, M. Haltmeier, and J. Schwab, “Deep learning for photoacoustic tomography from sparse data,” Inverse Problems in Science and Engineering 0, 1–19 (2018).

